# MARK2 in glial cells suppresses inflammatory responses and mitigates tau toxicity

**DOI:** 10.1101/2025.07.21.665902

**Authors:** Aoi Fukuchi, Sho Nakajima, Akiko Asada, Taro Saito, Kanae Ando

## Abstract

Neuroinflammation is a pathological hallmark of Alzheimer’s disease and related neurodegenerative diseases. However, signaling molecules that regulate glial activation status are not fully understood. Microtubule affinity-regulating kinase 2 (MARK2) has been implicated in both immune responses and AD pathology. Here, we report that MARK2 negatively regulates glial immune responses, which protects against neurodegeneration. We found that MARK2 knockdown in the BV2 murine microglial cell line enhanced IL-6 expression in response to LPS. MARK2 knockdown enhanced IL-6 expression induced by TLR7 agonist but not stimulation of RLR pathways and cGAS-STING. In the brains of PS19 tauopathy mice, MARK2 was elevated in homeostatic microglia but reduced in activated microglia. In *Drosophila* expressing human tau in the retina, expression of AMP downstream of the Toll pathway in the pigment glia enhances degeneration of photoreceptor neurons. Glial knockdown of Par-1, the Drosophila ortholog of MARK2, enhanced Toll-mediated AMP expression and neurodegeneration, whereas overexpression of Par-1 in the pigment glia suppressed them. These results suggest that MARK2/Par-1 in glia negatively regulates Toll pathway-driven inflammation and protects against tau-induced neurodegeneration. These findings provide insight into the molecular underpinnings of glial inflammation in neurodegenerative conditions and highlight MARK2 as a potential therapeutic target for modulating neuroinflammatory responses.

## Introduction

Neuroinflammation, a chronic inflammation in nervous tissue, is a common feature in neurodegenerative diseases (Heneka et al., 2015). Microglia are the resident macrophages in the CNS and regulate CNS development and maintenance via their immune functions against infections and injuries; however, microglia are overactivated and become destructive to neurons under disease conditions (Deczkowska et al., 2018)(Ransohoff, 2016; Streit, 2002). Such disease-associated microglia (DAM) can harm neurons via excess phagocytosis and the release of neurotoxic cytokines, contributing to neuron loss in disease pathogenesis (Chen and Holtzman, 2022). DAM are characterized molecularly with expression of microglial markers including *Iba1, Cst3*, and *Hexb* as well as downregulation of homeostatic microglial genes such as *P2ry12, P2ry13, Cx3cr1, CD33*, and *Tmem119*, and found in the patients brains with various neurodegenerative diseases such as Alzheimer’s disease and ALS as well as mouse models of them (Deczkowska et al., 2018). Glial activation in response to neurodegeneration is also observed in a Drosophila model of tauopathy (Oka et al., 2025), indicating that DAM is a common signature of microglial response to neurodegeneration.

Microglia produce proinflammatory cytokines, such as Interleukin-6 (IL-6), Interleukin-1 β (IL-1β), Tumor necrosis factor α (TNFα), Interferon α (IFNα), and Interferon β (IFNβ), under the control of innate immune sensors (Gao et al., 2023). Microglia utilize several immune sensors, including Toll-like receptors (TLRs), the cyclic GMP-AMP synthase (cGAS)-stimulator of interferon genes (STING) pathway, the RIG-I-like Receptor (RLR) pathway, inducing translocation of nuclear factor κB (NF-κB) to nuclei and cytokine transcription (Akinduro et al., 2025; Aravalli et al., 2007; Gern et al., 2024; Gulen et al., 2023; Huang et al., 2023). NF-κB-mediated innate immune response is conserved across species: *Drosophila* has two innate immune sensors, the Toll pathway activated by Gram-positive bacteria and by fungi, and the immune deficiency (Imd) pathway activated by Gram-negative bacterial infection, both of which activate NF-κB transcription factors to induce expression of antimicrobial peptides (AMPs) (Hetru and Hoffmann, 2009). Excess production of proinflammatory cytokines are detrimental, however, molecular mechanisms that modulate these signaling pathways to prevent overactivation are not fully understood.

Microtubule affinity-regulating kinase (MARK) 2 is a Ser/Thr protein kinase that belongs to an evolutionarily conserved Par-1 family, which regulates polarity and growth (Wu and Griffin, 2017). MARK2 is known to phosphorylate microtubule-associated protein tau in neurons (Drewes et al., 1997), which enhances tau neurotoxicity (Ballatore et al., 2007). MARK2 has also been associated with immune functions. Splicing variants of MARK2 are associated with inflammation (Hueso et al., 2004). MARK2-deficient mice show immune dysfunction and autoimmune diseases. Primary cultured microglial cells isolated from MARK2-deficient mice show altered differentiation and sensitivity to stimuli (Hurov et al., 2001). MARK2 regulates T cell polarity following engagement to an APC (Lin et al., 2009), modulates NF-κB transcripts in Caco-2 cells stimulated with cytokines (Mashukova et al., 2021), LPS-induced production of cytokines in macrophages (Deng et al., 2020), microglial morphological changes (DiBona et al., 2019), and UBAC2-mediated inflammatory responses (He et al., 2024), suggesting its critical role in the immune system in various tissues. However, it is not fully understood how microglial MARK2 affects neuroinflammation in neurodegenerative conditions. In this study, we analyzed the roles of MARK2 in microglial immune responses and the roles of glial MARK2 in tau toxicity using murine glial cell lines, a mouse model of tauopathy, and *a Drosophila* model of tauopathy. Our results suggest that MARK2 in glia negatively regulates cytokine expression and protects against neurodegeneration.

## Material and methods

### Mouse husbandry

PS19 was purchased from The Jackson Laboratories. All mice were housed in a special pathogen-free area at the animal housing facility of Tokyo Metropolitan University under a 12:12 h light/dark cycle, with ad libitum access to food (Picolab mouse diet 20, 5058) and water. This study was approved by the Research Ethics Committee of Tokyo Metropolitan University: A5-5, A5-6, A4-6, A4-23, A3-11. All animal experiments were performed in accordance with the Tokyo Metropolitan University Animal Experiment Guidelines and the guidelines of the Japan Neuroscience Society.

### Drosophila husbandry

Flies were reared on standard medium containing 10% glucose, 0.7% agar, 9% cornmeal, 4% brewer’s yeast, 0.3% propionic acid, and 0.1% butyl paraoxibenzoate [w/v] on a 12:12 h light/dark cycle at 25°C. Female flies 1∼2 or 7∼10 days old after eclosion were used in this study. UAS-Par-1 RNAi and UAS-Par-1Myc were kind gifts from Dr. Bingwei Lu (Stanford University) (ref). 54C-GAL4 was a gift from Dr. Shinya Yamamoto (Baylor College of Medicine) (ref). Gmr-hP301Ltau (Bloomington stock # 51377), Repo-GAL4 (Bloomington stock # 7415), UAS-luciferase (Bloomington stock # 35789), UAS-tdtomato (UAS-Cactus RNAi (NIG-fly stock #5858R-1) were obtained from NIG-fly (Japan) or the Bloomington stock center (Indiana University). UAS-luciferase RNAi has been described previously (Iijima-Ando et al., 2012). This study was approved by the Research Ethics Committee of Tokyo Metropolitan University: G5-16.

### Tissue culture

BV2 cells (Elabscience (Houston, TX)) were cultured in MEM (11095-080, Thermo Fisher Scientific) medium at 5% CO_2_ and 37°C. This study was approved by the Research Ethics Committee of Tokyo Metropolitan University: G5-16.

### LPS treatment

LPS was purchased from Sigma-Aldrich (L2654), 3p-hpRNA was purchased from Invivogen (tlrl-hprna-100), Imiquimod (14956) and DMXAA (14617) were purchased from Cayman. LPS and 3p-hpRNA were dissolved in UltraPure DNase/RNase-Free Distilled Water (10977-015, Invitrogen), and Imiquimod and DMXAA was dissolved in DMSO (046-21981, Fujifilm Wako Pure Chemical Corporation).

### RNA interference

MISSION siRNA Universal Negative Control #1 and MARK2 siRNA were purchased from Sigma-Aldrich. The sequences are as follows.

si-mouse MARK2 #1: 5’-GCACUUUAGAGCAAAUUAU-3’

Lipofectamine RNAiMAX Reagent (13778-150, Invitrogen) was then used to transfect siRNA into BV2 cells according to the manufacturer’s instructions. Four hours after transfection, the medium was replaced with new medium. 1 day later, cells were treated with LPS or Imiquimod, 3p-hpRNA, DMXAA, and collected for further analysis.

### Subcellular Fractionation

BV2 cells were seeded in 12-well plates, treated with siRNA against MARK2, followed by LPS (1ng/ml) for 30 min, and collected separately into nuclear and cytoplasmic fractions using the Nuclei ez prep nuclei isolation kit (NUC-101, SIGMA). Histone H3 was used as a nuclear marker and GAPDH as a cytoplasmic marker.

### SDS-PAGE, Phos-tag SDS-PAGE, Western blotting

BV2 cells were seeded into 12 wells and treated with siRNA against MARK2, followed by treatment with LPS (1 ng/ml) for 30 min. Samples were also seeded into 12 wells and treated with LPS (100 ng/ml) for 30 min. Cells were lysed in 1% SDS 50 mM NaF solution, sonicated and boiled at 95°C for 3 min, and mixed with an equal volume of Laemmli sample buffer (10 μg/ml leupeptin, 0.4 μM pefabloc, 10 mM β-glycerphosphate, 10 mM NaF, 10% (v/v) β-mercaptoethanol), boiled again at 95°C for 3 minutes. For *Drosophila* samples, more than 15 female heads were collected at 1 to 2 days after eclosion and homogenized in Laemmli sample buffer, boiled at 95°C for 2 min. Phos-tag SDS-PAGE was performed using a 7.5% (w/v) polyacrylamide gel containing 50 μM Phos-tag (AAL-107, Wako Chemicals), 100 μM MnCl_2_. Proteins were transferred to PVDF membrane (IPVH00010, Millipore) using a crown-water transfer device (XCell SureLock Mini-Cell XCell II Blot Module, EI0002, Invitrogen). The membrane was then blocked with 5% skim milk (TBS solution containing 0.05% Tween-20) or 5% BSA (TBS solution containing 0.05% Tween-20) for 1 hour at room temperature. Reacted with the following primary antibodies diluted in blocking solution or Can Get Signal solution (NKB-101, TOYOBO) for at least 12 hours at 4°C. Anti-MARK2 antibody (15492-1-AP, Proteintech), anti-Actin antibody (A2066, Sigma-Aldrich), anti-Nf-κB antibody (6956, Cell Signaling technology), anti-GAPDH antibody (NB100-56875, Novus Biologicals), anti-IkB antibody (4814, Cell Signaling technology), anti-pT595MARK2 (ab34751, abcam), anti-pT208 MARK2 (PA5-17495, Invitrogen), anti-Tau antibody (T46, 13-6400, Invitrogen), anti-pS262Tau antibody (ab131354, abcam) were purchased. Secondary antibodies, goat anti-mouse immunoglobulin HRP (P0447, Agilent), swine anti-rabbit immunoglobulin HRP (P0399, Agilent), were used and detected by Immobilon Western Chemiluminescent HRP Substrate (WBKLS0500, Merck Millipore). Chemiluminescence signals were observed using Fusion FX (Vilber) and quantified using ImageJ (NIH).

### RT-qPCR

BV2 cells were seeded in 12-well plates and treated with siRNA against MARK2, followed by 6 h LPS (1 ng/ml), Imiquimod (1 μg/ml), 3p-hpRNA (100 ng/ml), DMXAA (10 μg/ml). Then, total RNA was extracted using Isogen reagent (311-02501, Nippon Gene) and TE buffer (pH 8.0) (314-90021, Nippon Gene). For Drosophila, more than 30 female heads were frozen 1-2 days after eclosion, and total RNA was extracted using Isogen reagent (311-02501, Nippon Gene). Total RNA was reverse transcribed using ReverTra Ace qPCR RT Master Mix with gDNA Remover (FSQ-301, TOYOBO). Quantitative PCR was performed on a Thermal Cycler Dice Real Time System (TP-800, TAKARA Bio), with Cycle Value (CT) averages calculated using at least three replicates in each sample. Expression levels of genes of interest were standardized using GAPDH (glyceraldehyde-3-phosphate dehydrogenase), rp49 (ribosomal protein 49) or Actin. Relative expression levels were identified by the ΔΔCT method. Primers were designed using Primer-BLAST (NIH) or the DRSC Fly Primer Bank (Harvard Medical School).

Mouse IL-6 for 5’ -CTTGGGACTGATGCTGGGTGACA-3’

Mouse IL-6 rev 5’ -GCCTCCGACTTGTGAAGTGGTA-3’

Mouse IFNb for 5’ -CTGGAGCAGCTGAATGGAAAG-3’

Mouse IFNb rev 5’ -CTTCTCCGTCATCTCCATAGGG-3’

Mouse IL-1b for 5’ -TGACGGGACCCCAAAAGATGA-3’

Mouse IL-1b rev 5’ -TCTCCACAGCCACAATGAGT-3’

Mouse TNFa for 5’ -CCCTCACACTCAGATCATCTTCT-3’

Mouse TNFa rev 5’ -GCTACGACGTGGGCTACAG-3’

Mouse IL-10 for 5’ -GCTCTTACTGACTGGCATGAG-3’

Mouse IL-10 rev 5’ -CGCAGCTCTCTAGGAGCATGTG-3’

Mouse IFNa for 5’ -CTTCCACAGGATCACTGTGTACCT -3’

Mouse IFNa rev 5’ -TTCTGCTCTCTGACCACCTCCC-3’

Mouse GAPDH for 5’ -TGTGTGTCCGTCGTGGATCTGA -3’

Mouse GAPDH rev 5’ -TTGCTGTTGAAGTCGCAGGAG -3’

Drosophila Cactus for 5’ -ATGCCGAGCCCAACAAAAG -3’

Drosophila Cactus rev 5’ -CGCTAGTGGCTAGTGAGGAC -3’

Drosophila Par-1 for 5’ -CGCTGGAATACTCGGGGCAC -3’

Drosophila Par-1 rev 5’ -AGAAGAAGCTCATGCGACCC -3’

Drosophila Attacin C for 5’-CTGCACTGGACTACTCCCACATCA-3’

Drosophila Attacin C rev 5’ -CGATCCTGCGACTGCCAAAGATTG-3’

Drosophila Cecropin A1 for 5’ -CATTGGACAATCGGAAGCTGGGTG-3’

Drosophila Cecropin A1 rev 5’ -TAATCATCGTGGTCAACCTCGGGGC-3’

Drosophila Diptericin B for 5’ -AGGATTCGATCTGAGCCTCAACGGG-3’

Drosophila Diptericin B rev 5’ -TGAAGGTATACACTCCACCGGGCTC-3’

Drosophila Drosomycin for 5’ -AGTACTTGTTCGCCCTCTTCGCTG-3’

Drosophila Drosomycin rev 5’ -CCTTGTATCTTCCGGGACAGGCAGT-3’

Drosophila Metchnikowin for 5’ -CATCAATCAATTCCCGCCACCGAG-3’

Drosophila Metchnikowin rev 5’ -AAATGGGTCCCTGGGTGACGATGAG-3’

Drosophila rp49 for 5’ -GCTAAGCTGTCGCACAAATG-3’

Drosophila rp49 rev 5’ -GTTCGATCCGTAACCGATGT-3’

Drosophila actin 5C for 5’ -TGCACCGCAAGTGCTTCTAA-3’

Drosophila actin 5C rev 5’ -TGCTGCACTCCAAACTTCCA-3’

### Immunohistochemical Staining

Mice were deeply anesthetized with (8)double doses of 0.3 mg/kg medetomidine, 4.0 mg/kg midazolam, and 5.0 mg/kg butorphanol. Mice were then perfused with PBS, followed by 4% paraformaldehyde (PFA)/PBS. Mouse brains were then removed and fixed in 4% PFA/PBS solution for 24 hours. The brains were then immersed in 30% sucrose PBS until the brain sank to the bottom of a 15 ml Eppendorf tube. Frozen sections were then prepared at 40 µm thickness, and sections were immersed in cryoprotective solution (30% ethylene glycol, 20% glycerol, 40% ddH_2_O, 10% 10xPBS) and stored at - 28°C until use. Mouse brain sections were incubated with 5% normal goat serum (S-1000-20, vector laboratories) or normal donkey serum (017-000-121, Jackson ImmunoResearch) (selected by the host of secondary antibody) and 0.5% BSA (0.3% Triton X-100 in PBS solution) and the antibody was blocked with 3% normal goat serum or normal donkey serum and diluted with 0.5% BSA/PBST solution. Human brain sections were treated with deparaffinized, rehydrated, and autoclaved in TE buffer (pH 9.0) for antigen activation at 121°C for 20 min. Blocked with 5% normal goat serum or normal donkey serum/PBST solution, and the antibody was diluted with 3% normal goat serum or normal donkey serum/PBST solution. After blocking, sections were reacted with the following primary antibodies at 4°C for at least 12 hours. Rabbit anti-MARK2 antibody (15492-1-AP, Proteintech), goat anti-Iba1 antibody (011-27991, Fujifilm Wako Pure Chemical Corporation), goat anti-GFAP antibody (GTX89226, GeneTex), rat anti-P2RY 12 antibody (848001, Biolegend), then brain sections were reacted with the following secondary antibodies for 2 hours at room temperature, and nuclei were counterstained with DAPI/PBS. Donkey anti-rabbit IgG Alexa647 (A-31573, Invitrogen), donkey anti-goat IgG Alexa488 (ab150133, Abcam), goat anti-rabbit IgG Alexa488 (A-11008, Invitrogen), goat anti-rat IgG Alexa594 (112 585-003, Jackson ImmunoResearch). Samples were stored in the dark at 4°C. Samples were observed with a Keyence BZ-X710 fluorescence microscope.

### Histological Analysis of the fly retina

Female Drosophila heads from 7to 10 days after eclosion were fixed in Bouin’s Fixative solution (Picric acid, Formalin, Glacial acetic acid) for 48 hours at room temperature, then incubated in 50 mM Tris/150 mM NaCl After 24 hours of incubation in buffer, paraffin sections were prepared; 7 µm thick sections were prepared, stained with hematoxylin and eosin, and observed under a brightfield microscope. Images containing Drosophila lamina were taken with a Keyence microscope BZ-X700 (Keyence), and the area of the vacuole was measured with ImageJ (NIH). Heads of three or more flies (four or more hemispheres) were analyzed for each genotype.

### Statistical analyses

Microsoft Excel (Microsoft) or R (R Foundation for Statistical Computing, Vienna, Austria. URL http://www.R-project.org/) was used for statistical processing of the data. Student’s t-test, One-way ANOVA followed by Tukey HSD test, and Fisher’s Exact Test were used to determine significant differences. A difference was considered significant if the minimum significance level was p < 0.05. * P < 0.05, ** P < 0.01, *** P < 0.001, N.S. indicated no significant difference. Values are presented as mean ± SD or± SE.

## Results

### MARK2 inhibits the expression of inflammatory cytokines

We set out to analyze the roles of MARK2 in the expression of proinflammatory cytokines. The microglial BV2 cell line expresses inflammatory cytokines such as IL-1β, IL-6, IL-10, TNFα, and IFNβ in response to LPS stimulation (Henn et al., 2009)(Figure 1A). siRNA-mediated knockdown of MARK2 (Figure 1A) enhanced LPS-induced expression of IL-1β, IL-6, and TNFα significantly, while IFNβ induction was not altered (Figure 1 B). We also analyzed the expression of anti-inflammatory cytokines. IL-10 expression was increased by LPS stimulation; however, MARK2 knockdown did not affect its expression level (Figure 1B). Expression of IFNα was not increased by LPS treatment or by MARK2 knockdown (Figure 1B). These results indicate that knockdown of MARK2 enhances the induction of expression of inflammatory cytokines.

**Figure 1.**
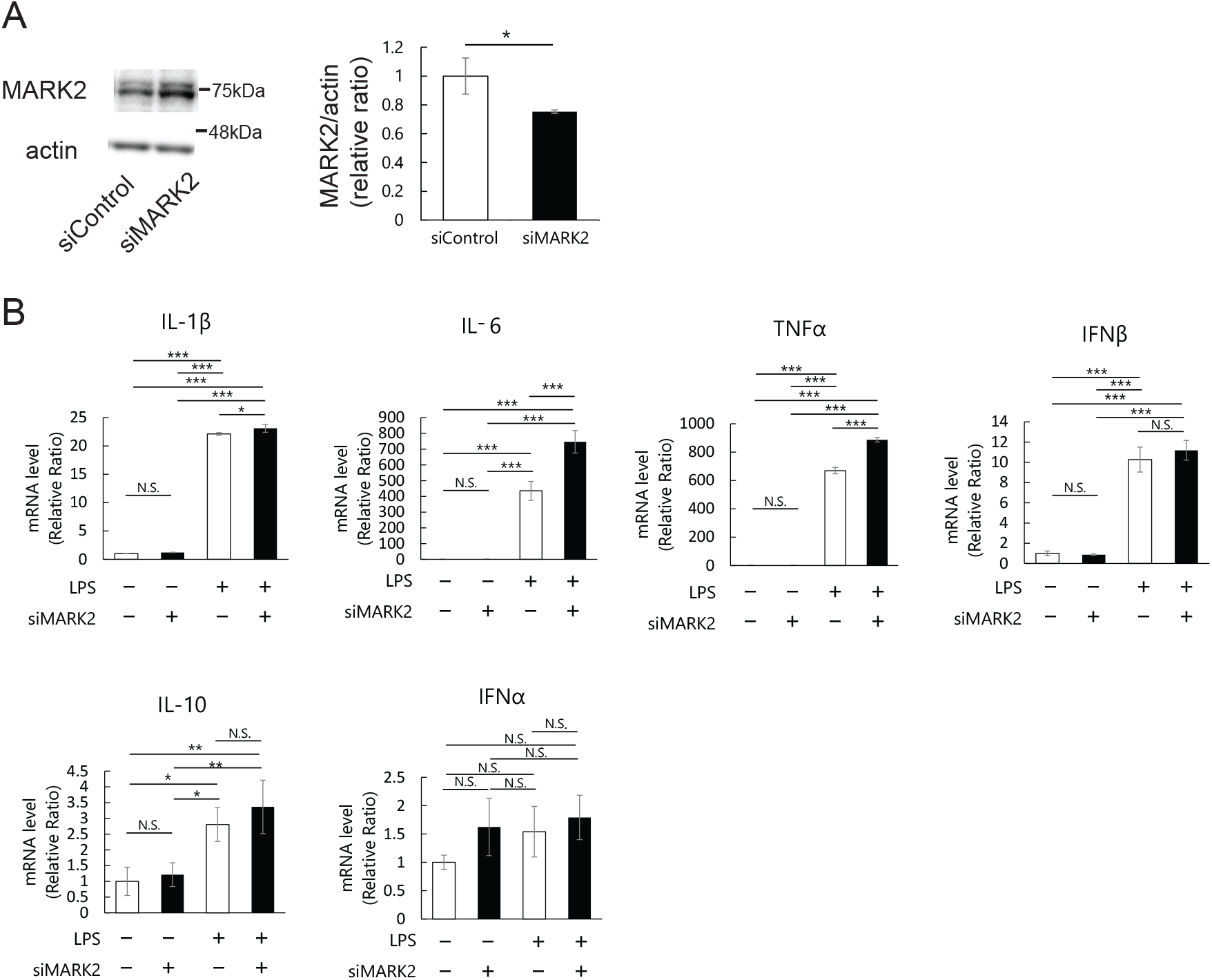
MARK2 knockdown enhances LPS-induced cytokine expression in BV2 cells. (A) MARK2 siRNA reduced MARK2 expression in BV2 cells. Western blot of BV2 cells treated with control siRNA or MARK2 siRNA. Actin was used as a loading control. Representative blots and quantitation are shown. N=3, mean ± _SD, *, p < 0.05 (Student’s t-test) (B) mRNA expression of IL-6, IFNβ, IL-1β, TNFα, IL-10, IFNα was analyzed via qRT-PCR. BV2 cells were transfected with control siRNA or MARK2 siRNA and treated with LPS (1ng/ml) for 6 hours. N=3, Mean ± SD, N.S., p>0.05, *, p<0.05, **, p<0.01, ***, p < 0.001 (One-way ANOVA followed by Tukey HSD test)

### MARK2 knockdown affects cytokine expression induced by TLR pathways

LPS activates TLR4 and induces cytokine expression (Lehnardt et al., 2002). We tested whether MARK2 affects cytokine expression induced by other pathways. TLR7, RLR pathways, and cGAS-STING were stimulated by imiquimod, 5’ triphosphate hairpin RNA (3p-hpRNA), and DMXAA, respectively. MARK2 knockdown enhanced cytokine expression induced by imiquimod (Figure 2A) but not those induced by 3p-hpRNA or DMXAA (Figure 2 B and C). These results suggest that MARK2 negatively regulates the expression of inflammatory cytokines through the TLR pathways.

**Figure 2.**
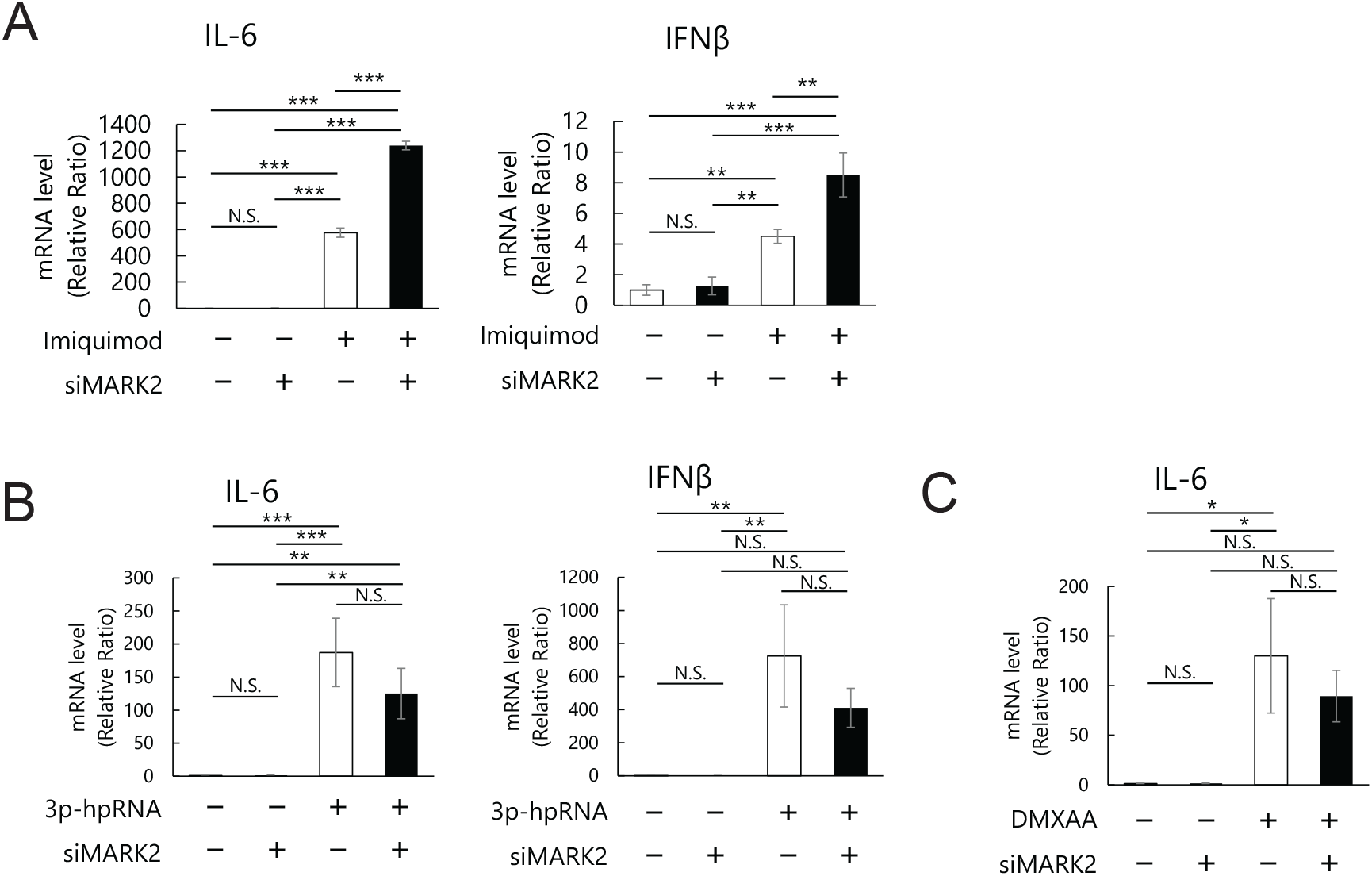
MARK2 negatively regulates IL-6 via the TLR pathway. (A-C) MARK2 knockdown enhances TLR-induced cytokine expression. BV2 cells were transfected with control or MARK2 siRNA and treated with Imiquimod (1μg/ml), 3p-hpRNA (100ng/ml), or DMXAA (10μg/ml) for 6 hours. Cells were subjected to qRT-PCR to analyze mRNA levels of IL-6 and IFNβ. N=3, Mean ± SD, N.S., p>0.05, **, p<0.01, ***, p < 0.001 (One-way ANOVA followed by Tukey HSD test).

### LPS treatment reduces MARK2 phosphorylation at Thr595

We asked whether activation of BV2 cells affects endogenous MARK2 activity. MARK2 activity is positively regulated by its phosphorylation at T208 by LKB1 (Lizcano et al., 2004) and negatively regulated by phosphorylation at T595 by aPKC (Hurov et al., 2004). We examined the phosphorylation levels of T208 and T595 of MARK2 by Western blotting. LPS stimulation did not affect total MARK2 levels or T208 phosphorylation and decreased T595 phosphorylation of MARK2 (Supplementary Figure 1). This result suggests that MARK2 activity is increased in response to LPS stimulation, which may be a part of a negative feedback mechanism to suppress excess activation.

### MARK2 expression is higher in homeostatic microglia and lower in activated microglia in the PS19 mouse brain

Results from BV2 cells suggest that MARK2 in microglia negatively regulates the production of inflammatory cytokines. If so, MARK2 expression may negatively correlate with glial activation status in the brain. In the brain of tauopathy model mice PS19, the number of homeostatic microglia is reduced, and that of activated microglia is increased (Deczkowska et al., 2018) (Supplementary figure 2). Co-immunostaining of MARK2 with homeostatic glial marker Purinergic Receptor P2Y12 (P2RY12) and that with activated microglial marker Iba1 revealed that MARK2 was expressed more often in P2RY12-positive microglia than in Iba1-positive cells (Figure 3). These results indicate that in the PS19 mouse model of tauopathy, MARK2 expression is high in homeostatic microglia and low in activated microglia.

**Figure 3.**
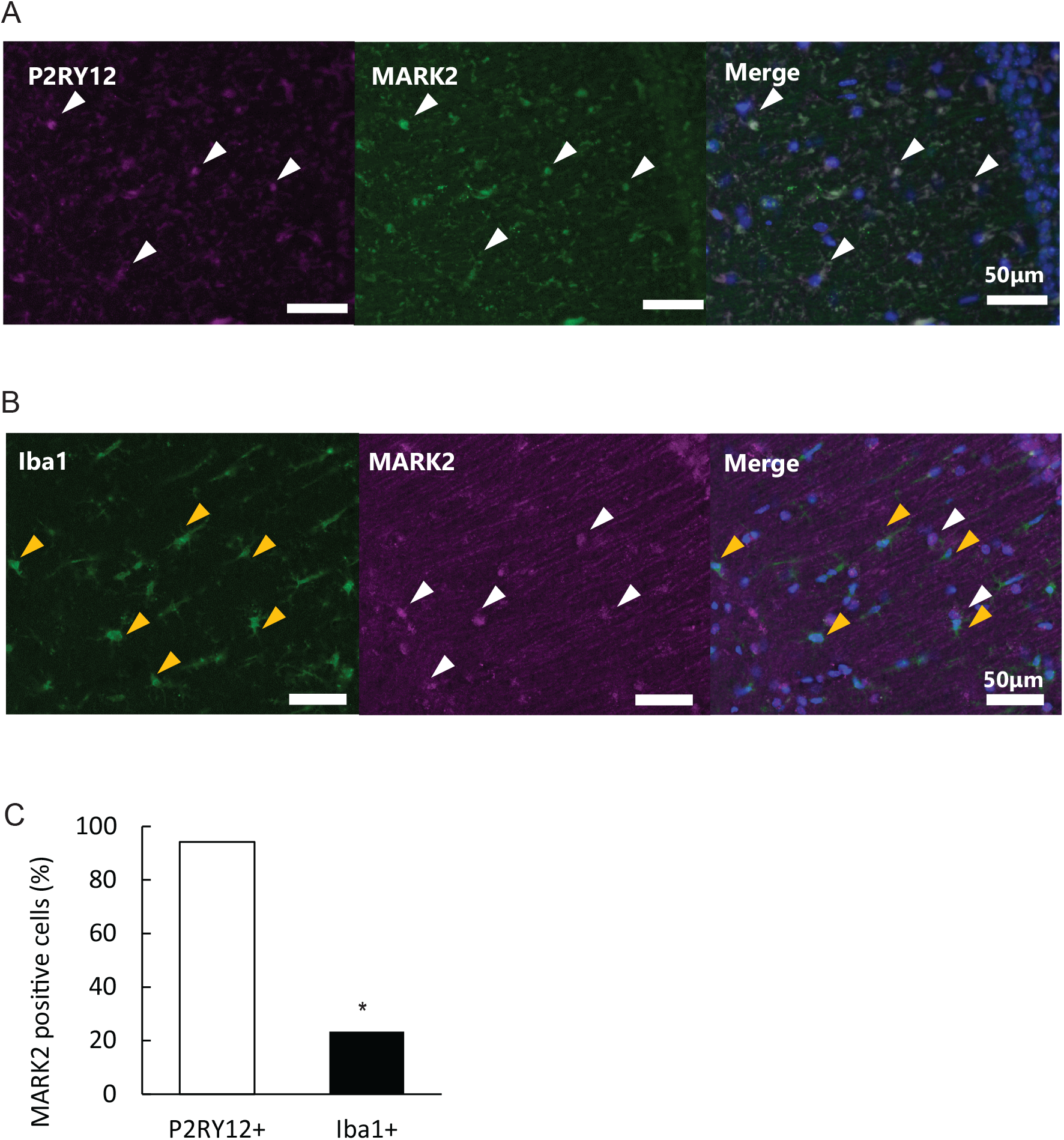
MARK2 is expressed in homeostatic microglia more often than in activated microglia. (A) Co-immunostaining of MARK2 and P2RY12 (top) or MARK2 and Iba1 (bottom) of PS19 brain. Sections are counterstained with DAPI. (Top) White arrows indicate MARK2-positive, P2RY12-positive cells. (Bottom) Yellow arrows in the bottom panels indicate MARK2-negative, Iba1-positive cells, and white arrows indicate MARK2-positive cells without Iba1 staining. (B) Percentages of MARK2-positive cells among P2RY12-positive cells or MARK2-positive cells among Iba1-positive cells in PS19 hippocampus. N=3, *, p<0.05, Fisher’s exact test.

### Glial activation in a tauopathy fly model is negatively regulated by *Par-1*

To investigate the roles of glial MARK2 in neurodegeneration, we used a *Drosophila* model of tauopathy. *Drosophila* express antimicrobial peptides (AMPs) as innate immunity effector molecules (Hetru and Hoffmann, 2009). Human tau expression in the retina causes degeneration of photoreceptor neurons along with expression of AMPs in the pigment glia (Oka et al., 2025). AMP expression is regulated by the IMD and the Toll pathways (Ferrandon et al., 2007). IMD pathway, homologous to the mammalian tumor necrosis factor receptor pathway and TRIF-dependent TLR pathway, activates Relish, whereas the Toll signaling pathway, which is similar to the interleukin-1 receptor (IL-1R) and MyD88-dependent TLR pathways, activates Dif (Ferrandon et al., 2007) (De Gregorio et al., 2002) (Figure 4A). Tau expression in the retina induces expression of *Drosomycin, Attacin, Cecropin, Diptelicin*, and *Mechinikowin* in the pigment glia (Oka et al., 2025). *Drosomycin* is induced via the Toll pathway, and *Attacin, Cecropin, Diptelicin*, and *Mechinikowin* are induced via the Imd pathway (Huang et al., 2010)(Figure 4A). We confirmed that knockdown of *cactus*, a negative regulator of *Dif*, in the pigment glia enhances expression of *Drosomycin* in tau flies (Figure 4B).

**Figure 4.**
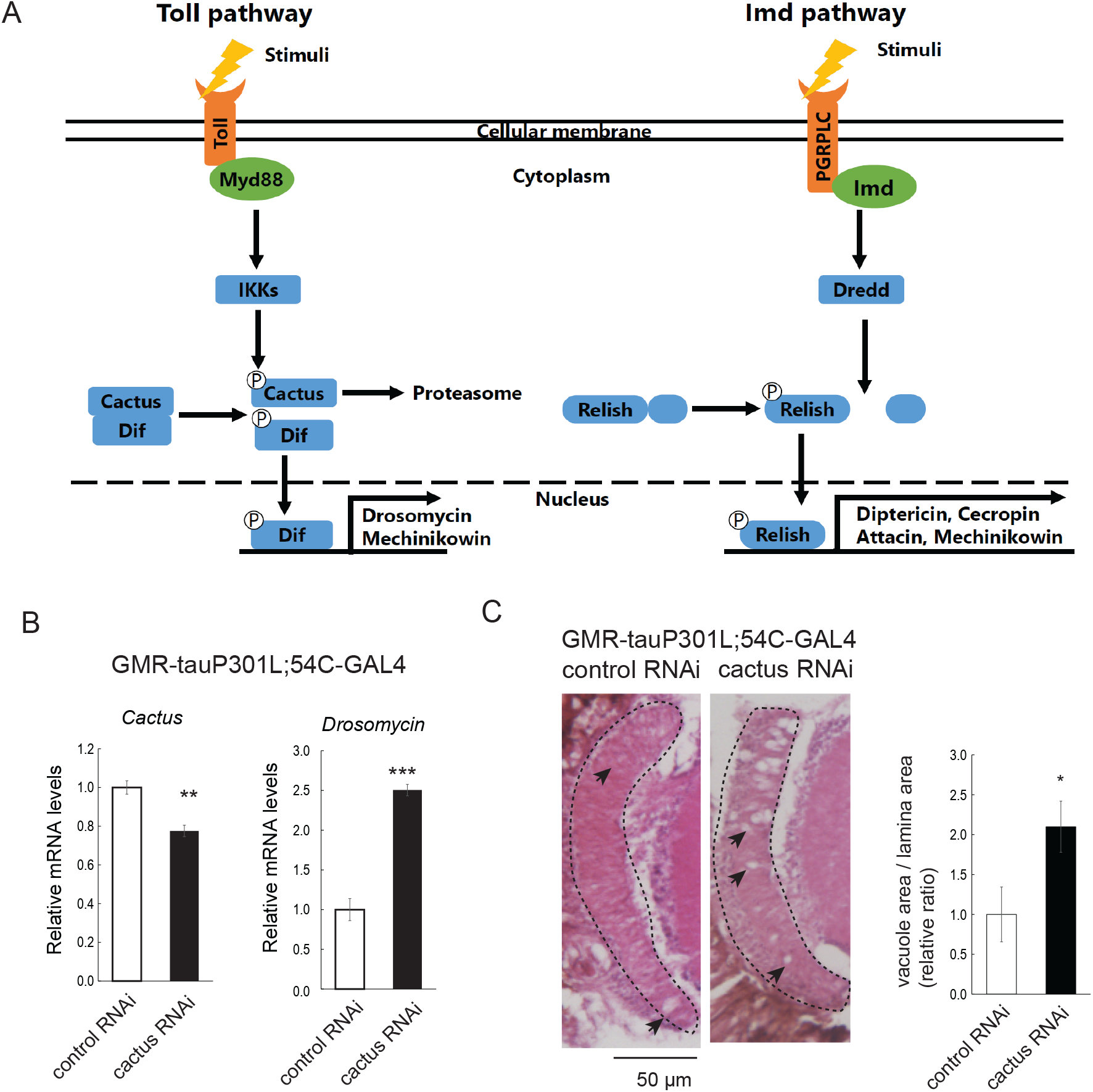
The Toll pathway in the pigment glia enhances tau-induced photoreceptor degeneration in a tauopathy fly model. (A) Schematic representation of Innate immune signaling pathways, Toll and IMD-NFkB, induce expression of AMPs in *Drosophila*. (B-C) cactus knockdown in the pigment glia enhances *Drosomycin* expression and photoreceptor degeneration in tau-expressing retina. (B) qRT-PCR of head extracts. N=3, mean ± SD, *, p < 0.05 (Student’s t-test). (C) H&E staining of retinal paraffin sections. Arrows indicate vacuoles. N=3, mean ± SD, *, p < 0.05 (Student’s t-test).

Tau expression in the retina induces degeneration of photoreceptor neurons during development, resulting in a smaller, deformed retina with a rough surface (rough-eye phenotype) (Wittmann et al., 2001) and age-dependent degeneration of photoreceptor axons, causing vacuoles in the lamina (Iijima-Ando et al., 2012). Cactus knockdown in glial cells enhanced tau-induced lamina degeneration, suggesting that activation of the *Toll* pathway enhances photoreceptor degeneration in the tau-expressing retina (Figure 4C).

We used this model to investigate how the *Drosophila* homolog of MARK2, *Par-1*, affects the expression of AMPs. Par-1 is the sole *Drosophila* homolog of MARK family members, and to analyze the function of Par-1 specifically in glia in the retina, we combined human tau expression under the direct control of *the GMR*-promoter with a tissue-specific expression system GAL4/UAS. To knock down *Par-1* specifically in glial cells, *Par-1* RNAi under UAS control was expressed with pigment, a glia-specific driver, 54C-GAL4. *Par-1* knockdown in the pigment glia in tau-expressing retina significantly increased expression of *Drosomycin* (Supplementary figure 3 and Figure 5A). Expression of Mechinikowin was not affected, and *Attacin, Cecropin*, and *Diptelicin* were significantly downregulated (Figure 5B). Similar results were obtained with knockdown of *Par-1* with a pan-glial driver, Repo-GAL4 (Figure 5B). In contrast, overexpression of Par-1 in the pigment glia reduced *Drosomycin* expression and increased expression of *Mechinikowin, Attacin, Cecropin*, and *Diptelicin* These results suggest that *Par-1* negatively regulate the *Toll*-*Dif*-*Drosomycin* axis in the pigment glia.

**Figure 5.**
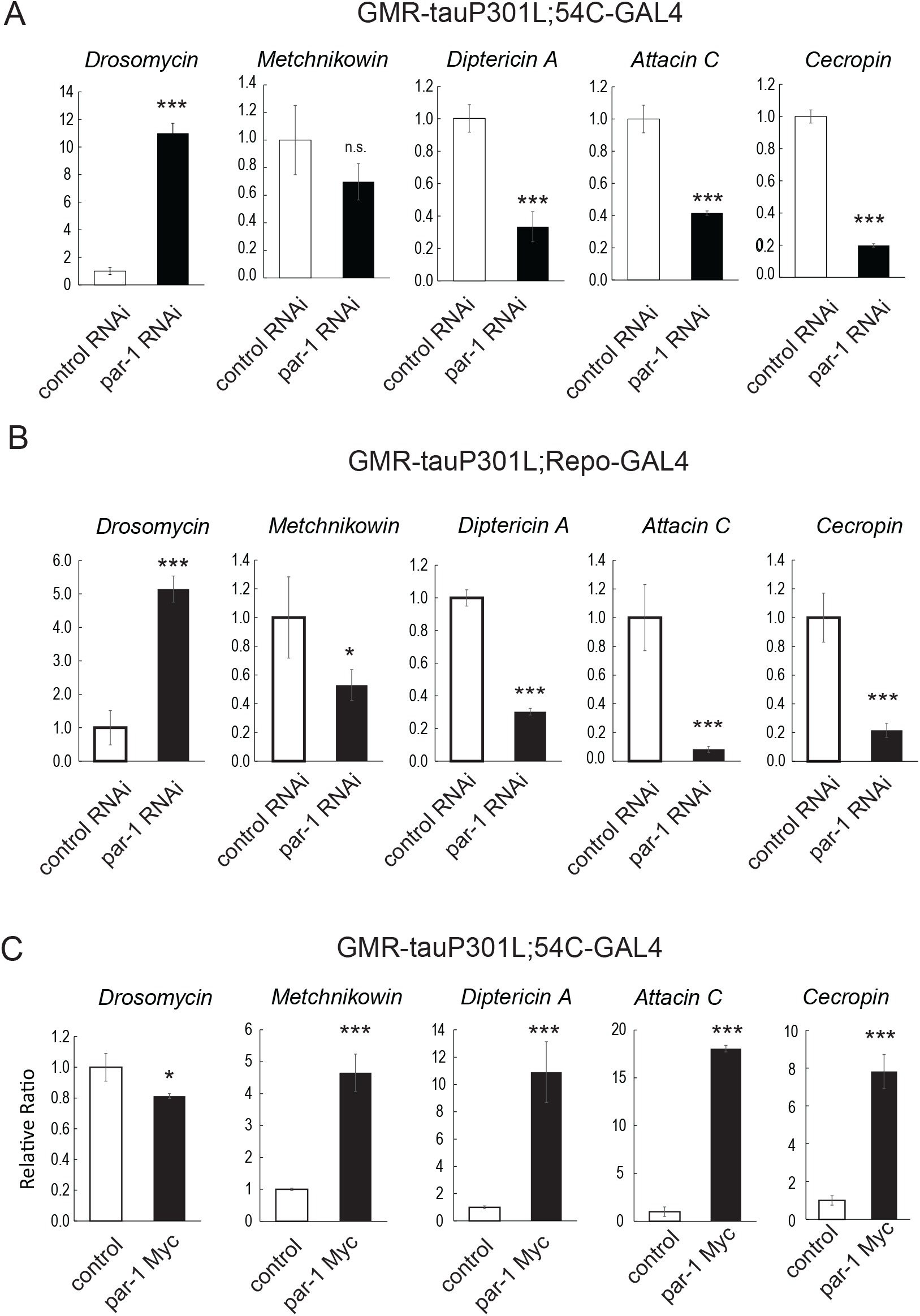
*Par-1* knockdown in the pigment glia increases *Drosomycin* expression. *Par-1* knockdown in pigment glia increases *Drosomycin* expression, but not *Metchinikowin* or *Attacin*. qRT-PCR of head extracts. Expression of Par-1 RNAi (A and B) or Par-1 (C) was driven by 54C-GAL4 (A and C) or Repo-GAL4 (B) in GMR-tauP301L background. More than 25 fly heads were pooled for each genotype. N=3, mean ± SD, *, p < 0.05 (Student’s t-test).

### Par-1 in the pigmented glia protects against tau-induced neurodegeneration

Finally, we examined the effects of *Par-1*-mediated glial immune responses on neurodegeneration in tau flies. Par-1 in the photoreceptor neurons is known to phosphorylate tau at Ser262 and Ser356, and promote tau-induced photoreceptor degeneration in a phosphorylation-dependent manner (Iijima et al., 2010; Iijima-Ando et al., 2012; Iijima-Ando et al., 2010). *Par-1* RNAi under UAS control was expressed by a pigment glia-specific driver 54C-GAL4 in the retina expressing tau under GMR promoter. Knockdown of *Par-1* in the pigment glia significantly increased photoreceptor degeneration (Figure 6A), and overexpression of Par-1 in the pigment glia mitigated photoreceptor degeneration (Figure 6 B). These results suggest that suppression of AMP expression by *Par-1* in pigment glia mitigates degeneration of photoreceptor neurons.

**Figure 6.**
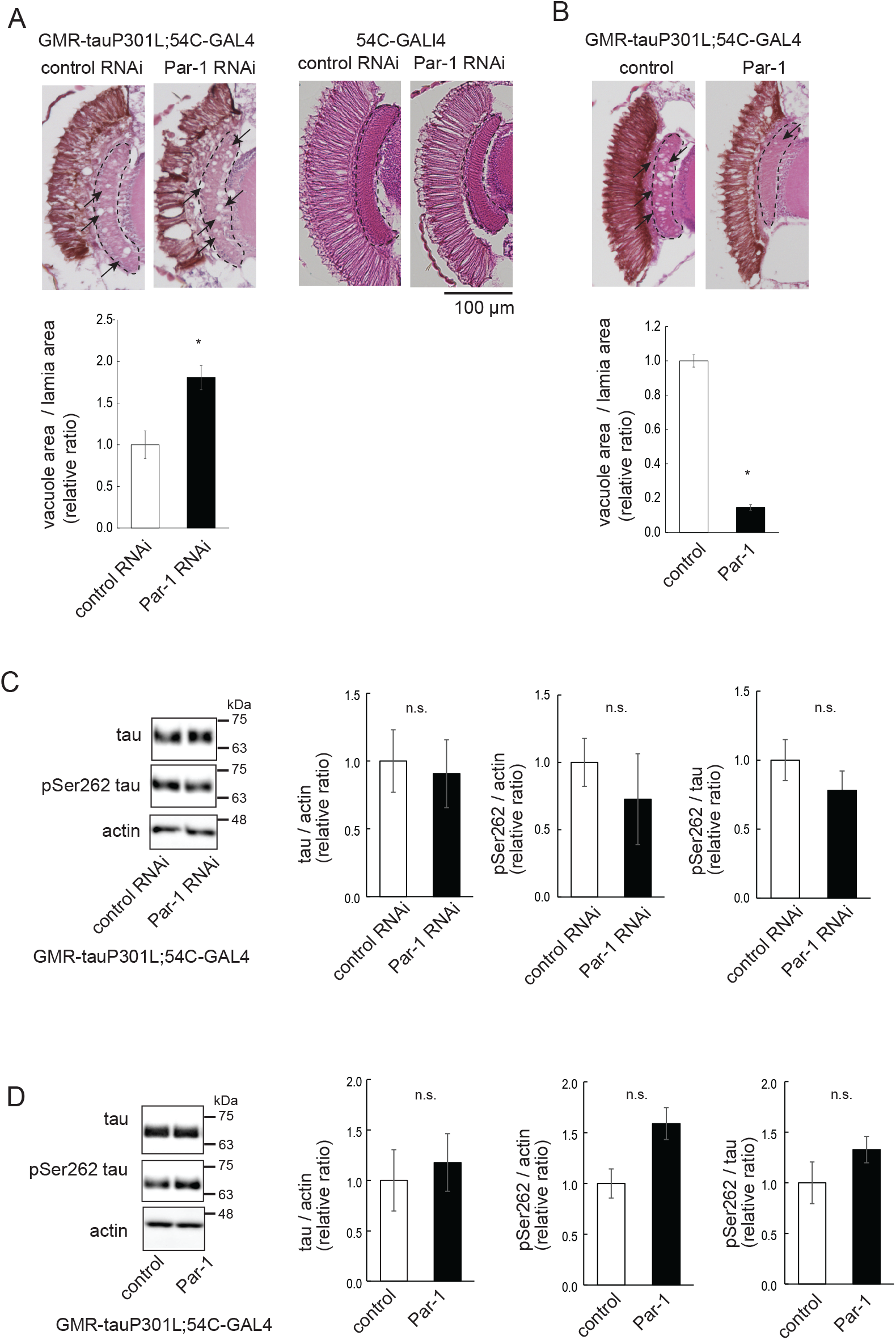
*Par-1* knockdown in the pigment glia enhances tau-induced neurodegeneration. H&E staining of retinal paraffin sections. Arrowheads indicate a vacuole. Representative images and quantitation of vacuole area. N=5, mean ± SE, *, p < 0.05 (Student’s t-test).

*Par-1* knockdown or overexpression in the pigment glia did not affect total tau levels in the retina (Figure 6C). Tau phosphorylated at Ser262 tends to decrease and increase with Par-1 knockdown or overexpression in the pigment glia, respectively; however, it did not reach statistical significance (Figure 6C). These results suggest that Par-1 in the pigment glia affects photoreceptor degeneration downstream of tau expression.

## Discussion

Neuroinflammation contributes to the disease progression of many neurodegenerative disorders. In this study, we found that MARK2 in glial cells plays a protective role against neuroinflammation. We found that MARK2 negatively regulates the expression of inflammatory cytokines induced by the TLR pathway in microglia-derived cultured cells. MARK2 was co-expressed with a homeostatic glial marker, P2RY12, more often than a marker of activated glia, Iba1. Glial-specific overexpression or knockdown in the Drosophila model of tauopathy revealed that Par-1 in the glial cells negatively regulates Drosomycin expression induced by the Toll pathway and protects photoreceptor neurons. These results suggest that MARK2 dysregulation may be involved in glial overactivation and the regulatory checkpoint of DAM under disease conditions.

We found that MARK2 negatively regulates the expression of a subset of proinflammatory cytokines in response to TLR4 activation (Figure 1). How MARK2 affects this signaling specifically is not clear at this point. After translocation to the nucleus, NF-kB proteins form different complexes with other transcriptional cofactors, which select subsets of the NF-kB– activated transcripts (Liu et al., 2017). Mashukova et al. reported that MARK2 phosphorylates Med17/TRAP80, a subunit of the Mediator complex, thus regulating NF-kB transcriptional functions in Caco-2 cells (Mashukova et al., 2021). In BV2 cells, MARK2 did not affect nuclear translocation of NF-kB after LPS stimulation or its phosphorylation (Supplementary Figure 4), suggesting MARK2 may also regulate transcription after these steps. In Caco-2 cells, MARK2 overexpression enhances TNFα responses (Mashukova et al., 2021), while our results suggest that MARK2 suppresses LPS-induced cytokine expression in BV2 cells (Figure 1). MARK2 may modulate NF-kB transcription in a cell-type or ligand-specific manner. MARK2 has also been reported to negatively regulate CBP (Tabassum et al., 2022), which can regulate NF-kB activity via acetylation (McKay and Cidlowski, 2000). Further analyses will be needed to elucidate the molecular details of MARK2-mediated regulation of the expression of inflammatory cytokines through the TLR pathways.

MARK2 and other MARK family members phosphorylate tau at Ser262 and Ser356 in the microtubule binding repeats (Drewes et al., 1997; Sultanakhmetov et al., 2024a). Tau phosphorylation at these sites enhances tau toxicity; thus, MARK2 is considered an intrinsic enhancer of tau toxicity. In fact, co-expression of tau and MARK2 in the entire retina increases tau phosphorylation at these sites and exacerbates photoreceptor degeneration in *Drosophila* (Sultanakhmetov et al., 2024a). *Par-1* in the photoreceptor neurons also promotes tau-induced photoreceptor degeneration in a Ser262/356-dependent manner (Chatterjee et al., 2009; Iijima-Ando et al., 2012; Iijima-Ando et al., 2010; Nishimura et al., 2004). In this study, we dissected the role of par-1 in the pigment glia by using the GAL4/UAS system in combination with tau under the direct control of *the GMR*-promoter. Knockdown or overexpression of *Par-1* specifically in the pigment glia in tau-expressing retina tends to decrease or increase Ser262 phosphorylation, respectively (Figure 6C); however, it did not reach statistical significance. Par-1 may phosphorylate tau in the pigment glia, but tau phosphorylation at these sites may mostly occur in photoreceptor neurons. Our results suggest that opposing roles of MARK2 in neurons and glia.

The mammalian Par-1/MARK family consists of MARK1-4. They are expressed in the brain and can phosphorylate tau proteins at Ser262 and Ser356. However, they have non-overlapping physiological functions, and they affect tau toxicity differently (Marx et al., 2010; Sultanakhmetov et al., 2024a; Wu and Griffin, 2017). MARK4 has the most significant enhancement among MARK1-4 in a fly model (Sultanakhmetov et al., 2024a). Their physiological functions are also different, and only MARK2 has been implicated in immune responses (Deng et al., 2020; DiBona et al., 2019; Hurov et al., 2001; Lin et al., 2009; Mashukova et al., 2021). MARK4 has been associated with Alzheimer’s disease genetically and pathologically (Lund et al., 2014; Pathak et al., 2020; Rovelet-Lecrux et al., 2015), and MARK4 deletion mitigates tau pathology and neuronal dysfunctions in PS19 mice (Sultanakhmetov et al., 2024b). Thus, MARK4 inhibition is considered a therapeutic strategy for Alzheimer’s disease. However, since kinase domains are highly conserved among MARK family members, competitive inhibitors currently available for MARK4 can also inhibit MARK2. We recently reported an allosteric MARK4-inhibitory peptide with high specificity (Oba et al., 2023), and non-competitive inhibitors may offer a promising avenue for drug development for MARKs.

## Abbreviations

MARK2: Microtubule affinity-regulating kinase 2
IL: interleukin
TLR: Toll-like receptor
NF-κB: The nuclear factor κB
IκB: cytosolic inhibitor of NF-κB
LPS: lipopolysaccharide
cGAS-STING: cyclic GMP-AMP synthase-stimulator of interferon genes
AMP: antimicroviral peptides

## Author contributions

Aoi Fukuchi: Conceptualization, Investigation, formal analyses, Writing – original draft

Sho Nakajima: Investigation, formal analyses

Akiko Asada: Supervision, Resources

Taro Saito: Conceptualization, Investigation, Resources, Supervision

Kanae Ando: Conceptualization, Funding acquisition, Investigation, Project administration,

Supervision, Writing – original draft, Writing – review & editing.

## Acknowledgments

Authors thank Sophia Limlingan (Tokyo Metropolitan University) and Grigorii Sultanakhmetov for technical help and comments, and Shin-Ichi Hisanaga and Dr. Adam Weitemier (Tokyo Metropolitan University) for critical comments.

## Data availability

The datasets used and/or analyzed in this study are available from the corresponding author upon request.

