## Supplementary Figure 1 for "MARK2 in glial cells suppresses inflammatory responses and mitigates tau toxicity"

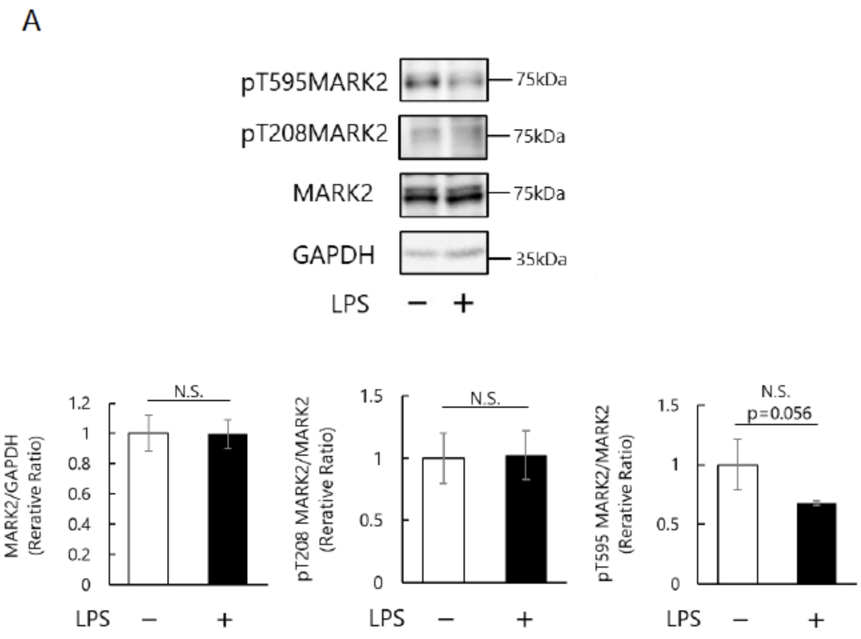


**Figure S1. MARK2 phosphorylation in BV2 cells after LPS stimulation.** BV2 cells were stimulated with LPS for 30 min and subjected to Western blotting with anti-MARK2 antibody, anti-pThr208 MARK2 antibody, or anti-pThr595 MARK2 antibody. GAPDH was used as a loading control. Representative blot (top) and quantitation (bottom). N = 3, Mean ± SD, N.S., $p>0.05$ (Student's t-test)
