## Supplementary Figure 2 for "MARK2 in glial cells suppresses inflammatory responses and mitigates tau toxicity"

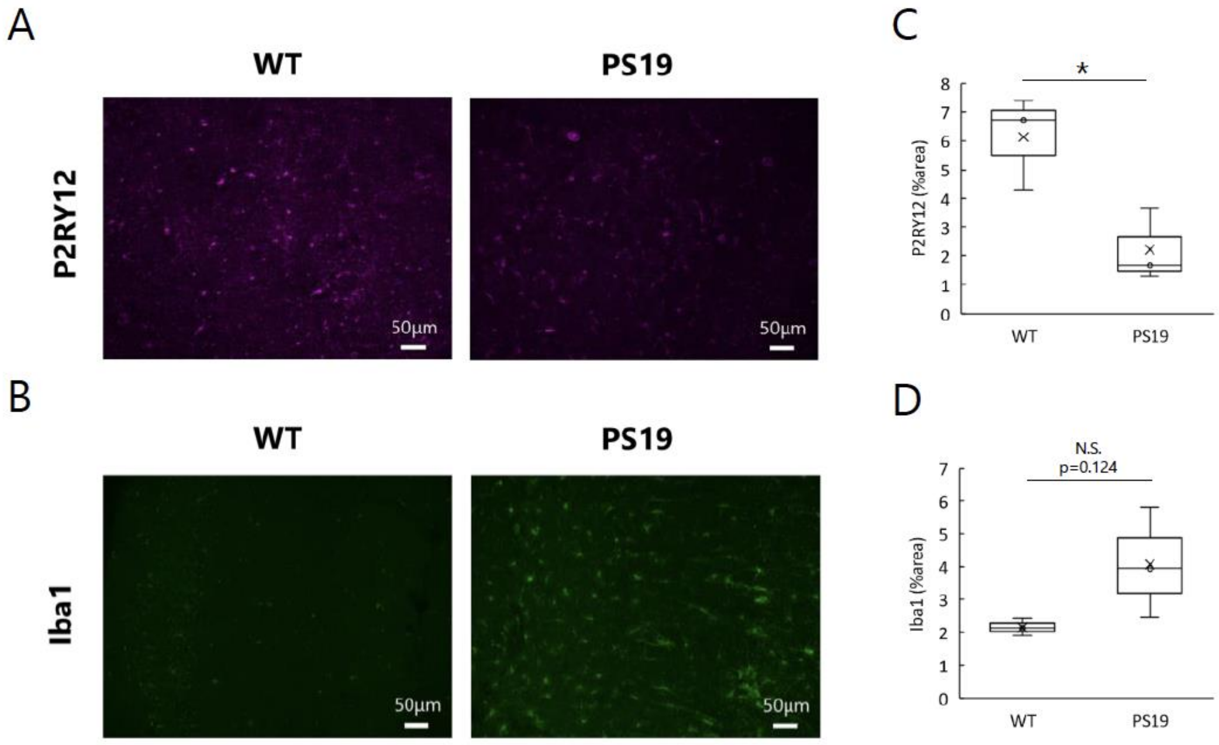


**Figure S2. The number of P2RY12-positive microglia reduces, and that of Iba1-positive microglia increases, in the tauopathy mouse model PS19 brain.**

Immunostaining of the hippocampus of 9-month-old control and PS19 male mice with anti-P2RY12 (A, C) and anti-Iba1 (B, D). (A, B) Representative images. Scale bar, 50 µm. (C, D) Quantitation of P2RY12-positive area and Iba-1-positive area. N=3, *p<0.05, Student's t-test.
