## Supplementary Figure 3 for "MARK2 in glial cells suppresses inflammatory responses and mitigates tau toxicity"

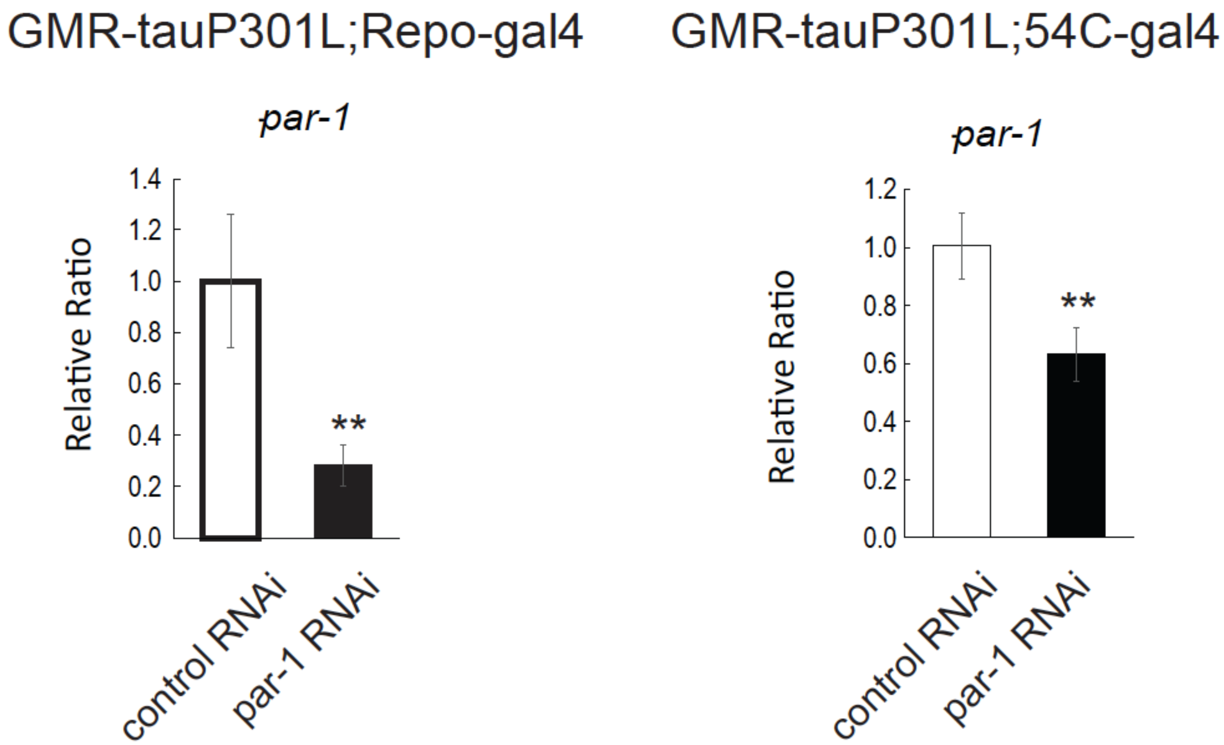


**Figure S3. Expression of Par-1 RNAi with pan-retinal driver Repo-Gal4 (Left) and the pigment glia-specific driver 54C-Gal4 (Right) lowers Par-1 expression.** qRT-PCR of head extracts. N=4, mean±SD, **, $p < 0.01$ (Student's t-test).
