## Supplementary Figure 4 for "MARK2 in glial cells suppresses inflammatory responses and mitigates tau toxicity"

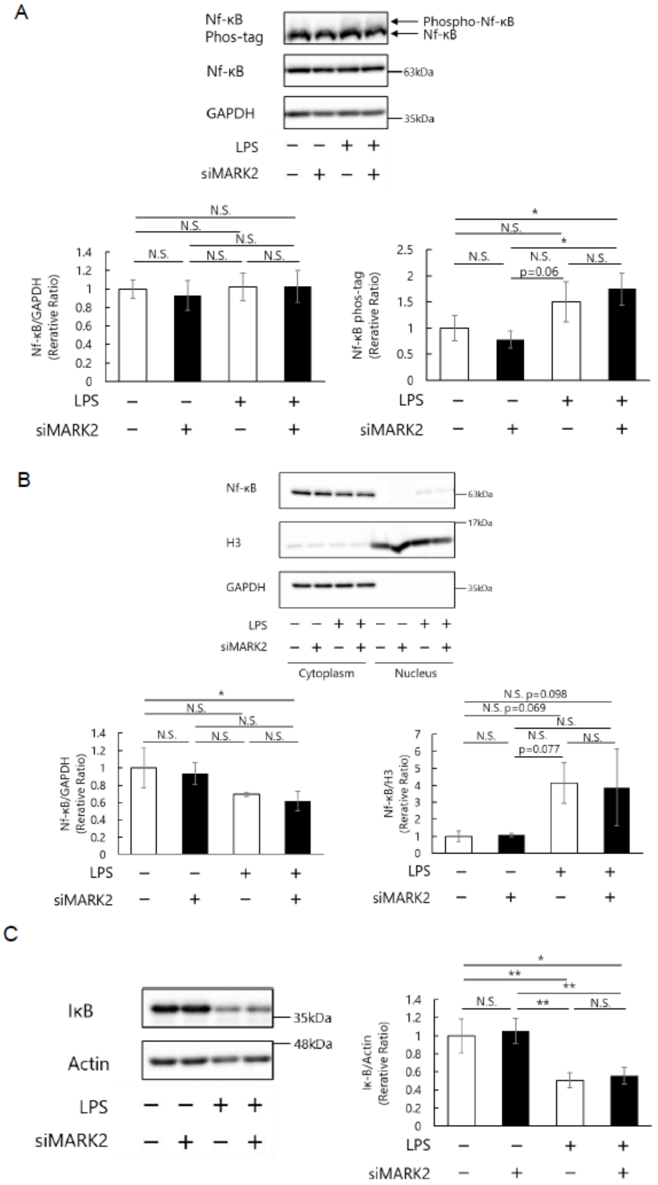


**Figure S4. MARK2 does not affect the nuclear translocation of Nf-κB**

(A) Phos-tag analyses of Nf-κB in BV2 cell lysate. GAPDH was used as a loading control. Representative blots and quantitation are shown. N＝3, Mean ± SD, N.S., p>0.05, *, p<0.05, One-way ANOVA followed by Tukey HSD test. (B) Western blot of Nf-κB in the cytosol and nuclear fractions of BV2 cells 30 min after LPS treatment. GAPDH and histone H3 were used as loading controls. N＝3, Mean ± SD, N.S., p>0.05, *, p<0.05, One-way ANOVA followed by Tukey HSD test. (C) Western blot of IκB. Actin was used as a loading control. N＝3, Mean± SD, N.S., p>0.05, *, p<0.05, **,p<0.01, One-way ANOVA followed by Tukey HSD test.
